# Functional variants in hematopoietic transcription factor footprints and their roles in the risk of immune system diseases

**DOI:** 10.1101/2021.03.22.436360

**Authors:** Naoto Kubota, Mikita Suyama

## Abstract

Genome-wide association studies (GWAS) have been performed to identify thousands of variants in the human genome as disease risk markers, but functional variants that actually affect gene regulation and their genomic features remain largely unknown. Here we performed a comprehensive survey of functional variants in the regulatory elements of the human genome. We integrated hematopoietic transcription factor (TF) footprints datasets generated by ENCODE project with multiple quantitative trait locus (QTL) datasets (eQTL, caQTL, bQTL, and hQTL) and investigated the associations of functional variants and immune system disease risk. We identified candidate regulatory variants highly linked with GWAS lead variants and found that they were strongly enriched in active enhancers in hematopoietic cells, emphasizing the clinical relevance of enhancers in disease risk. Moreover, we found some strong relationships between traits and hematopoietic cell types or TFs. We highlighted some credible regulatory variants and found that a variant, rs2291668, which potentially functions in the molecular pathogenesis of multiple sclerosis, is located within a TF footprint present in a protein-coding exon of the *TNFSF14* gene, indicating that protein-coding exons as well as noncoding regions can possess clinically relevant regulatory elements. Collectively, our results shed light on the molecular pathogenesis of immune system diseases. The methods described in this study can readily be applied to the study of the risk factors of other diseases.

## Introduction

Many individuals worldwide have complex diseases and traits caused by the combined effects of numerous genetic and environmental factors. Genome-wide association studies (GWASs) have been performed to identify the susceptibility regions of complex diseases in the human genome.^1^ However, for most complex diseases, the underlying genetic architecture and molecular details are still poorly understood. Most of the risk variants identified by GWASs occur in noncoding regions, so the discovery of causal variants and target genes has been challenging. Thus, the precise identification of the genes and pathways responsible for these complex diseases, which are possible diagnostic markers and therapeutic targets, is essential for the development of personalized genomic medicine.

Most of the causal variants of disease risk are predicted to affect the control of gene expression via regulatory elements such as promoters and enhancers.^2^ Variants that function in gene regulation, known as regulatory variants, alter the genomic sequence and can affect transcription factor (TF) binding levels, leading to the modulation of chromatin accessibility, histone modification levels, and gene expression. Datasets of genomic variants that correlate with quantitative traits, known as QTL, are useful for identifying regulatory variants.^3^ Recently, various types of QTL have been investigated, such as gene expression QTL (eQTL),^4–8^ chromatin accessibility QTL (caQTL),^9^ TF binding QTL (bQTL),^10,11^ and histone modification QTL (hQTL).^12^ Reanalysis and integration of multiple QTL can help identify credible functional variants. The prediction of TF occupancy has been improved by digital genomic footprinting, which is a computational approach that can identify TF binding sites within native chromatin at nucleotide resolution using data from chromatin accessibility assays such as DNase I-seq and ATAC-seq.^13–15^ The susceptibility regions of complex diseases have been identified at low resolution (∼kb scale), so most of the true causal variants remain unknown. Thus, identification of the precise position of causal variants and their roles in gene regulation is required to better understand the molecular mechanisms of disease risk.

In this study, we performed an integrated analysis of TF footprints and multiple QTL datasets from blood samples to identify overlooked functional variants affecting TF binding and regulatory activity and uncover more detailed mechanisms of disease risk. We focused on a comprehensive set of immune system diseases, including common autoimmune diseases, and developed a computational framework that enabled us to survey functional variants and prioritize candidate causal variants in 47 hematopoietic cell types. We examined the genomic and epigenomic features of candidate regulatory variants and investigated whether the variants associated with the risk of each trait are enriched in specific cell types and TFs. Based on this analysis, we found novel functional variants that may have been overlooked in previous studies and shed light on the molecular pathogenesis of immune system diseases.

## Material and Methods

### Identification of candidate regulatory variants

From the GWAS Catalog,^16^ we obtained summary data of GWAS lead variants and variants reportedly associated with immune system diseases, 238 reported traits (EFO ID: EFO_0000540), in the European population were used for the following analyses. We used LDlink^17^ to obtain data about variants that were highly linked (*r*^2^ > 0.8) with the GWAS lead variants in the European population. The genotype data in this database originated from Phase 3 (Version 5) of the 1000 Genomes Project, a public catalog of human variation and genotype data. Populations represented in the catalog include Utah Residents from North and West Europe, Toscani in Italia, Finnish in Finland, British in England and Scotland, and Iberian population in Spain.^18^ We obtained TF footprint datasets from the Encyclopedia of DNA Elements (ENCODE)^15^ for 47 biosamples, with a “Taxonomy_system” of “Hematopoietic,” and used them to identify variants located on TF binding sites using BEDTools.^19^ To find variants associated with differences in gene expression levels, we collected hematopoietic eQTL datasets from GTEx Analysis Release V8 (Cells - EBV-transformed lymphocytes and Whole Blood),^4^ GEUVADIS (lymphoblastoid cell lines),^5^ eQTLGen (whole blood),^6^ BLUEPRINT (CD14+ monocytes, CD16+ neutrophils, and naïve CD4+ T cells),^7^ and DICE (Activated naïve CD4+ T cells, Activated naïve CD8+ T cells, Classical monocytes, Memory T_REG_ cells, NK cells, Naïve B cells, Naïve CD4+ T cells, Naïve CD8+ T cells, Naïve T_REG_ cells, Non-classical monocytes, T_FH_ cells, T_H_1 cells, T_H_1/17 cells, T_H_17 cells, and T_H_2 cells).^8^ For subsequent analysis to identify more credible functional variants, we collected fine-mapped caQTL data;^9^ bQTL data for JUND, NFκB, POU2F1, PU-1, STAT1,^10^ and CTCF;^11^ and hQTL data for H3K27ac, H3K4me1, and H3K4me3,^12^ from lymphoblastoid cell lines.

### Genomic and epigenomic annotation

We used the annotatePeaks.pl function in HOMER (version 4.9.1)^20^ for annotation of the variants called multiple QTL (xQTL) to the following genomic elements: promoter, 5′ UTR, exon, intron, 3′ UTR, transcription termination site, intergenic, micro RNA, noncoding RNA, pseudogene, small cytoplasmic RNA, and small nucleolar RNA. We calculated the distance between each variant and the nearest TSS using the HOMER output files. To examine whether the eQTL variants (*n* = 5,565) were enriched in specific genomic elements, we used data produced by the chromatin state discovery and characterization software ChromHMM^21^ about the chromatin states of 127 biosamples, including 30 hematopoietic cells and immune tissues: E029 (primary monocytes from peripheral blood), E030 (primary neutrophils from peripheral blood), E031 (primary B cells from cord blood), E032 (primary B cells from peripheral blood), E033 (primary T cells from cord blood), E034 (primary T cells from peripheral blood), E035 (primary hematopoietic stem cells), E036 (primary hematopoietic stem cells short term culture), E037 (primary T helper memory cells from peripheral blood 2), E038 (primary T helper naïve cells from peripheral blood), E039 (primary T helper naïve cells from peripheral blood), E040 (primary T helper memory cells from peripheral blood 1), E041 (primary T helper cells PMA-I stimulated), E042 (primary T helper 17 cells PMA-I stimulated), E043 (primary T helper cells from peripheral blood), E044 (primary T regulatory cells from peripheral blood), E045 (primary T cells effector/memory enriched from peripheral blood), E046 (primary natural killer cells from peripheral blood), E047 (primary T killer naïve cells from peripheral blood), E048 (primary T killer memory cells from peripheral blood), E050 (primary hematopoietic stem cells G-CSF-mobilized female), E051 (primary hematopoietic stem cells G-CSF-mobilized male), E062 (primary mononuclear cells from peripheral blood), E093 (fetal thymus), E112 (thymus), E113 (spleen), E115 (Dnd41 T cell leukemia cell line), E116 (GM12878 lymphoblastoid cell line), E123 (K562 leukemia cell line), and E124 (monocytes-CD14+ RO01746 cell line). The chromatin states fall into 25 categories: 1_TssA (Active TSS), 2_PromU (Promoter Upstream TSS), 3_PromD1 (Promoter Downstream TSS 1), 4_PromD2 (Promoter Downstream TSS 2), 5_Tx5’’ (Transcribed -5’ preferential) 6_Tx (Strong transcription), 7_Tx3’’ (Transcribed - 3’ preferential), 8_TxWk (Weak transcription), 9_TxReg (Transcribed & regulatory (Prom/Enh)), 10_TxEnh5’’ (Transcribed 5’ preferential and Enh), 11_TxEnh3’’ (Transcribed 3’ preferential and Enh), 12_TxEnhW (Transcribed and Weak Enhancer), 13_EnhA1 (Active Enhancer 1), 14_EnhA2 (Active Enhancer 2), 15_EnhAF (Active Enhancer Flank), 16_EnhW1 (Weak Enhancer 1), 17_EnhW2 (Weak Enhancer 2), 18_EnhAc (Primary H3K27ac possible Enhancer), 19_DNase (Primary DNase), 20_ZNF/Rpts (ZNF genes & repeats), 21_Het (Heterochromatin), 22_PromP (Poised Promoter), 23_PromBiv (Bivalent Promoter), 24_ReprPC (Repressed Polycomb), and 25_Quies (Quiescent/Low). For each state, we computed the log odds ratio of eQTL variants in comparison to hematopoietic TF footprints, which is the merged region of TF footprints for 47 hematopoietic biosamples. We used the Mann–Whitney U test to compare the log odds ratios between hematopoietic cells/immune tissues (*n* = 30) and others (*n* = 97) for each state. The *P-*values were adjusted based on the false discovery rate using the Benjamini– Hochberg method.^22^ *P*-values of <0.05 were considered statistically significant.

### Enrichment analysis

Enrichment analysis was performed on 17 traits: allergic disease (asthma, hay fever, or eczema), ankylosing spondylitis, celiac disease, chronic lymphocytic leukemia (CLL), Crohn’s disease, hay fever and/or eczema, inflammatory bowel disease, multiple myeloma (MM), multiple sclerosis, psoriasis, rheumatoid arthritis, rheumatoid arthritis (ACPA-positive), systemic lupus erythematosus, systemic sclerosis, type 1 diabetes, ulcerative colitis, and vitiligo. To analyze the enrichment in the 47 hematopoietic cell types (trait–cell), we computed the enrichment score for each trait using the following procedure: we counted the number of eQTL variants present within the TF footprints for each cell type, normalized them by the total length of footprints and total number of LD-expanded risk variants, and finally transformed the values to *Z*-scores in each trait. We performed hierarchical clustering (average-linkage with euclidean metric) based on the enrichment score. We then downloaded “hub_186875_JASPAR2020_TFBS_hg38,” a dataset in which binding motifs in hg38 were defined using the JASPAR CORE database,^23^ from the UCSC Table Browser in bed format and identified motifs with track scores of >400. To analyze the enrichment in 722 TF types (trait–TF), we counted the number of eQTL variants present on TF binding motifs within hematopoietic TF footprints. Next, we computed the log odds ratio of eQTL variants for each TF in comparison with the entire genome. As well as trait–cell enrichment analysis, we performed hierarchical clustering based on the log odds ratio for 38 TFs, in which the log odds ratio was >8 for at least one trait.

### Functional analysis of candidate regulatory variants

To visualize the genomic positions around the candidate regulatory variants, we used the UCSC Genome Browser.^24^ The logos of TF binding motifs overlapping with candidate regulatory variants were downloaded from the JASPAR CORE database. To examine the allelic imbalance of DNase I-seq reads, we used processed data from the ENCODE project using beta-binomial distribution,^15^ and *P*-values of <0.05 were considered statistically significant. The plots of the effects of candidate regulatory variants on gene expression were obtained from the GTEx portal. Statistical significance was ascertained by the GTEx project (https://www.gtexportal.org/home/documentationPage).

### Statistical tests, visualization, and tools used

The computational framework used in this work was developed in Python 3.7 and GNU Bash 3.2 environment, and all statistical tests were conducted in this framework. All graphs, except for the GTEx violin plots, were constructed using the seaborn Python package.

## Results

### A computational framework for discovering regulatory variants involved in the risk of immune system diseases

We developed a computational framework to discover regulatory variants associated with immune system disease risk (Figure 1A). First, we obtained data on the variants that are considered markers of immune system diseases, such as autoimmune diseases and lymphoma (238 reported traits, which are registered in The Experimental Factor Ontology as EFO_0000540), in the European population from the GWAS Catalog (*n* = 3,954) (Table S1). Then, to survey potentially functional variants, we expanded variants with high linkage disequilibrium (*r*^2^ > 0.8 in the European population, 1000 Genomes) (*n* = 73,668). From these, the variants present within hematopoietic TF footprints from 47 cell types were obtained (*n* = 6,591) and used in the subsequent analysis. Next, we filtered the variants using multi QTL datasets of whole blood or immune cells, including gene expression levels (eQTL), chromatin accessibility (caQTL), TF binding levels (bQTL), and histone modifications (hQTL). For eQTL, we collected several datasets from GTEx, GEUVADIS, eQTLGen, BLUEPRINT, and DICE. We also used data about the variants showing allelic imbalance in DNase I-seq reads from ENCODE as caQTL in this framework. We hypothesized that the variants called multiple QTL (xQTL) could be more credible and clinically relevant. Regulatory variants should affect gene expression levels; therefore, the dataset was filtered to yield only variants called eQTL (*n* = 5,565). The eQTL variants overlapped with other types of QTL, such as caQTL, bQTL, and hQTL (Figure 1B, Figure S1, Table S2). We found that 35.3% of all eQTL variants (1,967/5,565) were also called other types of QTL. Sixty-seven variants were positive for all QTL (e+ca+b+h), indicating probable regulatory functions.

**Figure 1.**
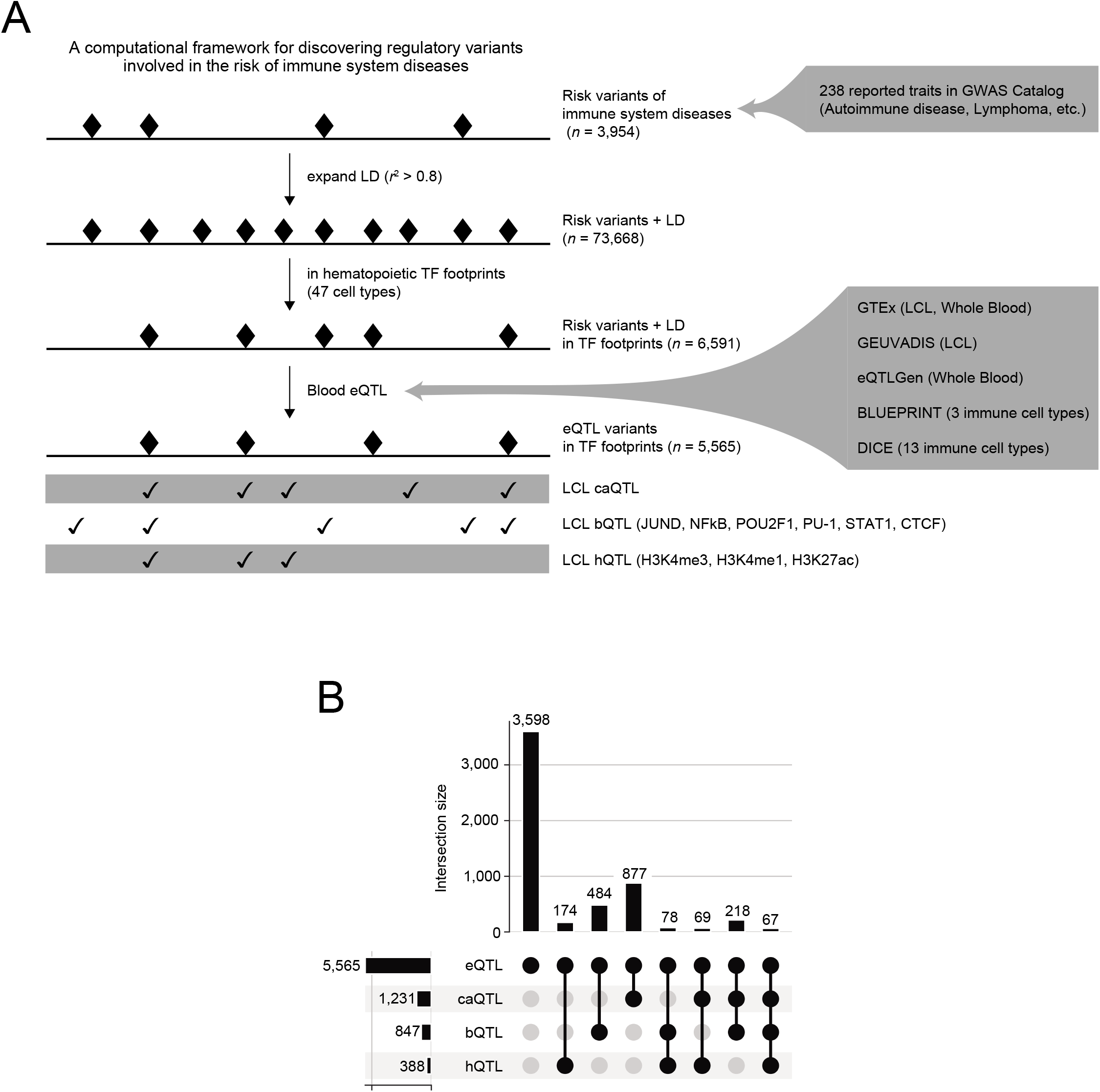
An overview of this study. (A) Schematic diagram of a computational framework to identify variants predicted to have regulatory functions. (B) The number of variants (y-axis) for each combination (columns) of QTL (rows).

### Genomic and epigenomic features of xQTL variants

We tested the genomic distribution of each xQTL variant group and found that all groups were enriched primarily in promoters and 5′ UTR regions (Figure 2A), in concordance with our understanding of the way in which variants at TFBS can affect gene expression. They can modulate promoter activities and gene expression levels through changes in chromatin accessibility, histone modification levels, and TF binding levels. The xQTL variants tended to be close to TSS, but some groups, including the all-positive group (e+ca+b+h), showed weak enrichment in TSS (Figure 2B). This observation suggests that credible functional variants are distributed not only in TSS-proximal regions but also in distal regulatory elements, including enhancers. The fraction of variants present within promoters (−2000 bp and +2000 bp from the TSS) was at most 40% (Figure 2C), suggesting that most functional variants were present in distal regulatory elements.

**Figure 2.**
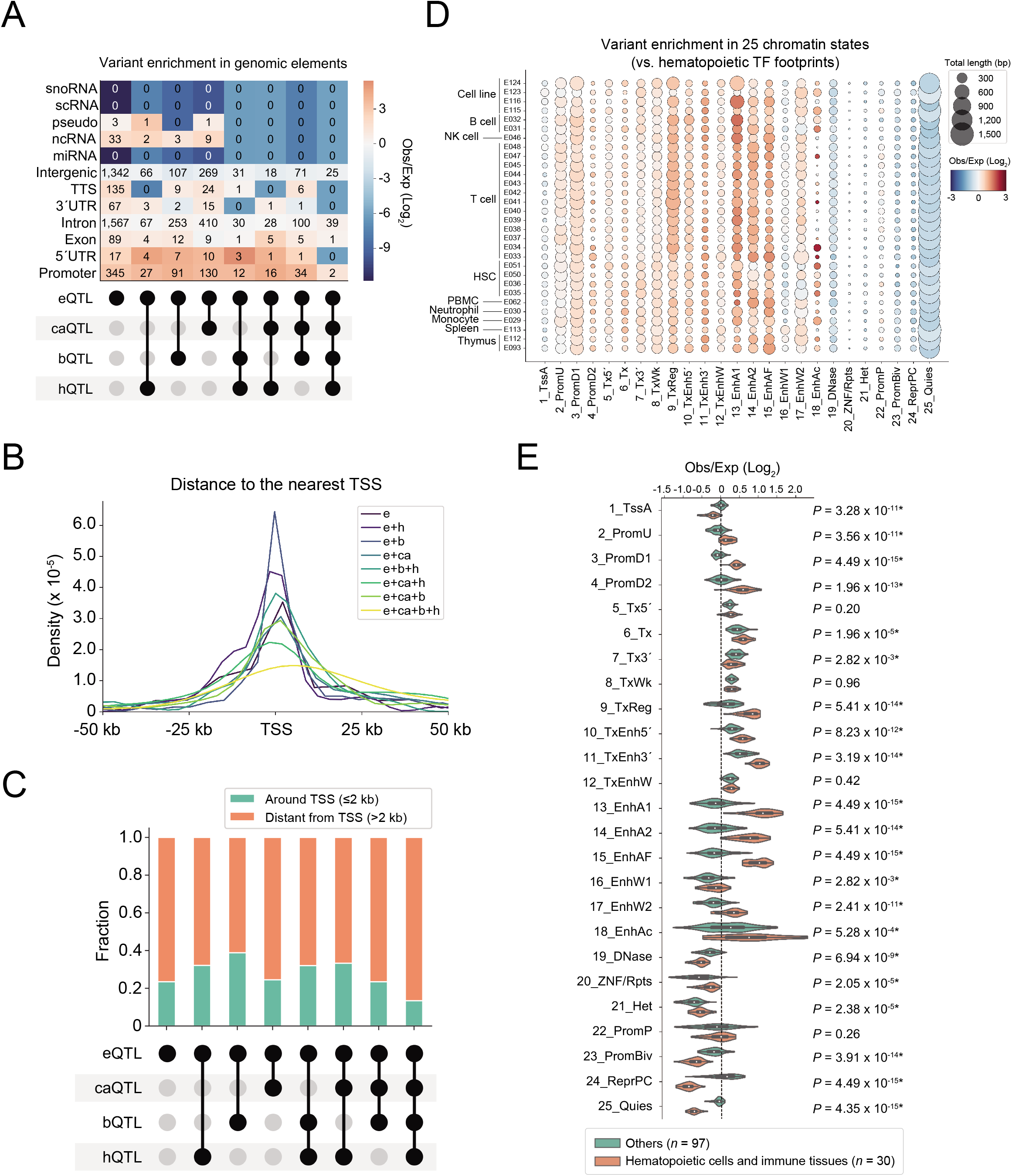
Genomic and epigenomic features of xQTL variants. (A) Enrichment in genomic elements for each combination of QTL. The numbers in each box represent the size of variants assigned to each element, and the color indicates the log odds ratio. (B) Distribution of distances between xQTL variants and their nearest TSS. (C) Fraction of xQTL variants located around (green) and distal (orange) to the nearest TSS. (D) Enrichment of eQTL variants (*n* = 5,565) in 25 chromatin states defined by ChromHMM^21^ for hematopoietic cells and immune tissues (*n* = 30). The circle size and color represent the total length of variants and the log odds ratio, respectively. (E) Comparison of the log odds ratio between hematopoietic cells/immune tissues (orange) and others (green). Adjusted *P*-values calculated using the Mann–Whitney U tests for each state are shown. The asterisks indicate statistical significance (*P* < 0.05).

Next, we used epigenomic annotation data defined by ChromHMM to estimate the log odds of eQTL variants to one of the 25 chromatin states in comparison to hematopoietic TF footprints. In blood cells and immune tissues, eQTL variants were strongly enriched in enhancer states (13_EnhA1, 14_EnhA2, 15_EnhAF, and 18_EnhAc) rather than in active TSS and promoter states (1_TssA, 2_PromU, 3_PromD1, and 4_PromD2), especially in B cells and T cells (Figure 2D). Compared with other cell types, eQTL variants were more strongly enriched in enhancer states (13_EnhA1, 14_EnhA2, 15_EnhAF, and 18_EnhAc) in hematopoietic cells and immune tissues (Figure 2E). Moreover, eQTL variants were less enriched in the active TSS state (1_TssA) in hematopoietic cells and immune tissues. These results emphasize that enhancers are actually relevant for understanding the genetic architecture of immune system diseases.

### Disease-linked variant enrichment in hematopoietic cells and TFs

Disease-linked variants are expected to be enriched in the regulatory sequences of specific cell types because each trait involves specific pathological pathways. We therefore analyzed the enrichment of eQTL variants linked with 17 major traits in the TF footprints of 47 hematopoietic cell types (Figure 3A). As expected, most autoimmune diseases were enriched in the TF footprints of T cells (h.CD8+-DS17203, h.CD4+-DS17881, h.CD3+-DS17198, hTreg-DS14702, etc.) and B cells (CD19+-DS17186 and CD20+-DS18208) but not in CD34+ hematopoietic stem cells (HSCs) (h.CD34+-DS12274, h.CD34+-DS12734, etc.). We found that variants linked with the risk of psoriasis were most strongly enriched in the TF footprints of CD4+ type 1 helper T cells (hTH1-DS7840), in agreement with previous findings.^25,26^ In contrast, the variants linked with the risk of Crohn’s disease were more strongly enriched in the TF footprints of CD8+ killer T cells (h.CD8+-DS17203) and CD56+ NK cells (h.CD56+-DS17189) rather than CD4+ helper T cells (h.CD4+-DS17881). Interestingly, some autoimmune diseases showed distinctive patterns of enrichment. For example, variants linked with the risk of type 1 diabetes were strongly enriched in the TF footprints of NAMALWA and Karpas, which are lymphoma-derived cell lines (h.NAMALWA-DS22816, h.NAMALWA-DS22823, h.Karpas-422-DS27641, and h.Karpas-422-DS27648), and those linked with the risk of vitiligo were strongly enriched in some CD34+ HSCs, which are supposed to be well-differentiated to erythroblasts (h.CD34+-DS24190, h.CD34+-DS24222, and h.CD34+-DS25969).^27^ The mechanisms underlying these associations between diseases and cell types are unclear, but these results may explain the difference in immunopathogenesis from other autoimmune diseases. Unlike most autoimmune diseases, variants linked with the risk of MM, a form of blood cancer, were strongly enriched in the TF footprints of an MM-derived cell line (h.b.lymphoblast.mm.1s-DS27522). This result indicates that MM-related variants are preferably present within the TF binding sites that are active in the blood cells of patients with MM, suggesting that they have functional importance in MM progression and not only in the onset of the disease. In CLL, another type of blood cancer, the variants linked with risk were strongly enriched in the TF footprints of CD34+ HSCs (h.CD34+-DS24190, h.CD34+-DS24222, and h.CD34+-DS25969) in addition to B cells (CD19+-DS17186). CLL is believed to originate from mature lymphocytes, but a previous study has reported that HSCs are also involved in the pathogenesis of the disease.^28^ This result might reflect the role of HSCs in CLL pathogenesis. Collectively, these results suggest that variant enrichment analysis in hematopoietic TF footprints can reveal their specific pathologically relevant cells.

**Figure 3.**
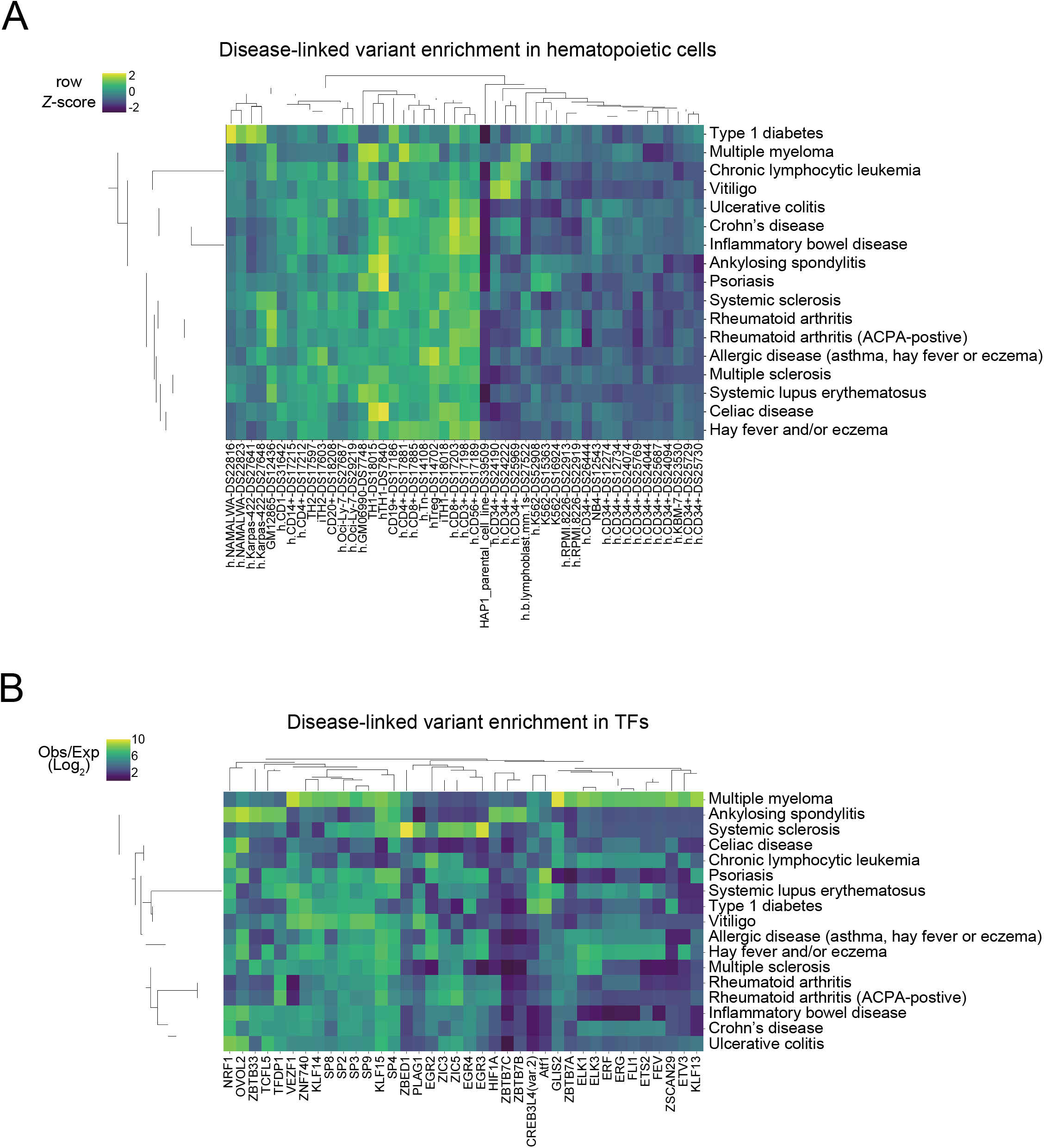
Disease-linked variant enrichment in hematopoietic cell types and TFs. (A) Hierarchically clustered trait–cell enrichment. The y-axis shows traits and the x-axis shows cell IDs of ENCODE TF footprints. The color indicates row *Z*-scores. (B) Hierarchically clustered trait–TF enrichment. The y-axis shows traits and the x-axis shows TF types. The color indicates log odds ratio.

We also performed enrichment analysis for trait–TF associations. The log odds of eQTL variants to one of 722 TF binding motif types in comparison to the whole genome were estimated according to trait. The result of this estimation for 12,274 pairs (17 traits × 722 TFs) revealed over-represented combinations such as systemic sclerosis and ZBED1 (log odds = 10.46) (Figure S2). Clustering of traits using log odds revealed differences in TF regulation among traits (Figure 3B). We found that MM had a distinct enrichment pattern. The variants linked with MM were strongly enriched in binding motifs of ETS factors (ELK1, ELK3, ERF, ERG, FLI1, ETS2, FEV, and ETV3). Consistent with our findings, a recent study reported that ETS factor binding motifs were significantly enriched in active enhancers of MM cells,^29^ suggesting the clinical relevance of ETS factors as key TFs in the risk of MM. We also found that MM-related variants were strongly enriched in the GLIS2 binding motif. The *CBFA2T3*-*GLIS2* fusion transcript was repeatedly identified in patients with acute myeloid leukemia (AML) and was associated with a poor prognosis.^30,31^ A recent study reported that Glis2 promotes differentiation of AML stem cells in mice.^32^ These results suggest that a change in GLIS2 regulation induces abnormal cell proliferation, leading to a risk of MM. We also found that the variants linked with the risks of most traits, specifically systemic sclerosis, were strongly enriched in binding motifs of EGR factors (EGR2, EGR3, and EGR4). These observations corroborate those of previous studies that have reported that deletion of both Egr2 and Egr3 in lymphocytes resulted in a systemic autoimmune response in mice.^33,34^ The strongest trait–TF combination in this enrichment analysis was the link between systemic sclerosis and ZBED1 (Figure S2). ZBED1 is a human homolog of the *Drosophila* DNA replication-related element-binding factor (DREF) and plays a pivotal role in cell proliferation.^35,36^ Its function in the immune system and autoimmune responses remains unclear, but ZBED1 might be a novel key TF in the risk of systemic sclerosis. Overall, our results suggest the existence of differences in TF regulation among diseases, resulting in distinctive molecular pathogenesis.

### New mechanistic insights into candidate regulatory variants

From the analysis performed using our computational framework, we identified candidate regulatory variants affecting gene expression, chromatin accessibility, and histone modifications. One variant identified, rs11748158 (C/T), was highly linked with rs13361189 (C/T), a GWAS lead variant of Crohn’s disease^37^ (*r*^2^ = 0.99), and with rs11749391 (C/T), a pleiotropic GWAS lead variant of chronic inflammatory diseases^38^ (*r*^2^ = 0.98) (Figure 4A). This regulatory variant was located within the helper and killer T cell-specific TF footprint (h.CD4+-DS17212 and h.CD8+-DS17885) within the 5′ UTR region of the *IRGM* gene. The chromatin state of this region was classified as promoter upstream TSS (2_PromU). Its minor allele (rs11748158-T, allele frequency = 0.1054 in the European population, 1000 Genomes) was predicted to result in decreased binding of KLF17 (Figure 4B) and to alter chromatin accessibility and the levels of H3K27ac, H3K4me1, and H3K4me3 (Figure 4C), indicative of active regulatory elements. Using data on DNase I-seq allelic imbalance,^15^ significantly fewer rs11748158-T allelic reads were mapped (*P* = 0.036, beta-binomial test) in rs11748158 heterozygous samples (Figure 4D), indicating that rs11748158-T decreases the regulatory activity. We confirmed that *IRGM* expression was significantly decreased depending on the rs11748158-T dosage in whole blood samples in the GTEx eQTL datasets (*P* = 2.7 × 10^−5^) (Figure 4E). A previous study reported that IRGM negatively regulates the inflammatory response by controlling inflammasome assembly.^39^ Consistent with our results, rs11748158 has been reported, using a deep learning-based framework, ExPecto, to be a putative causal variant of the risk of Crohn’s disease.^40^ The predicted effect of the variant on *IRGM* expression was functionally validated. Thus, our results reinforce the findings of Zhou et al. by providing a mechanistic interpretation based on currently available public datasets.

**Figure 4.**
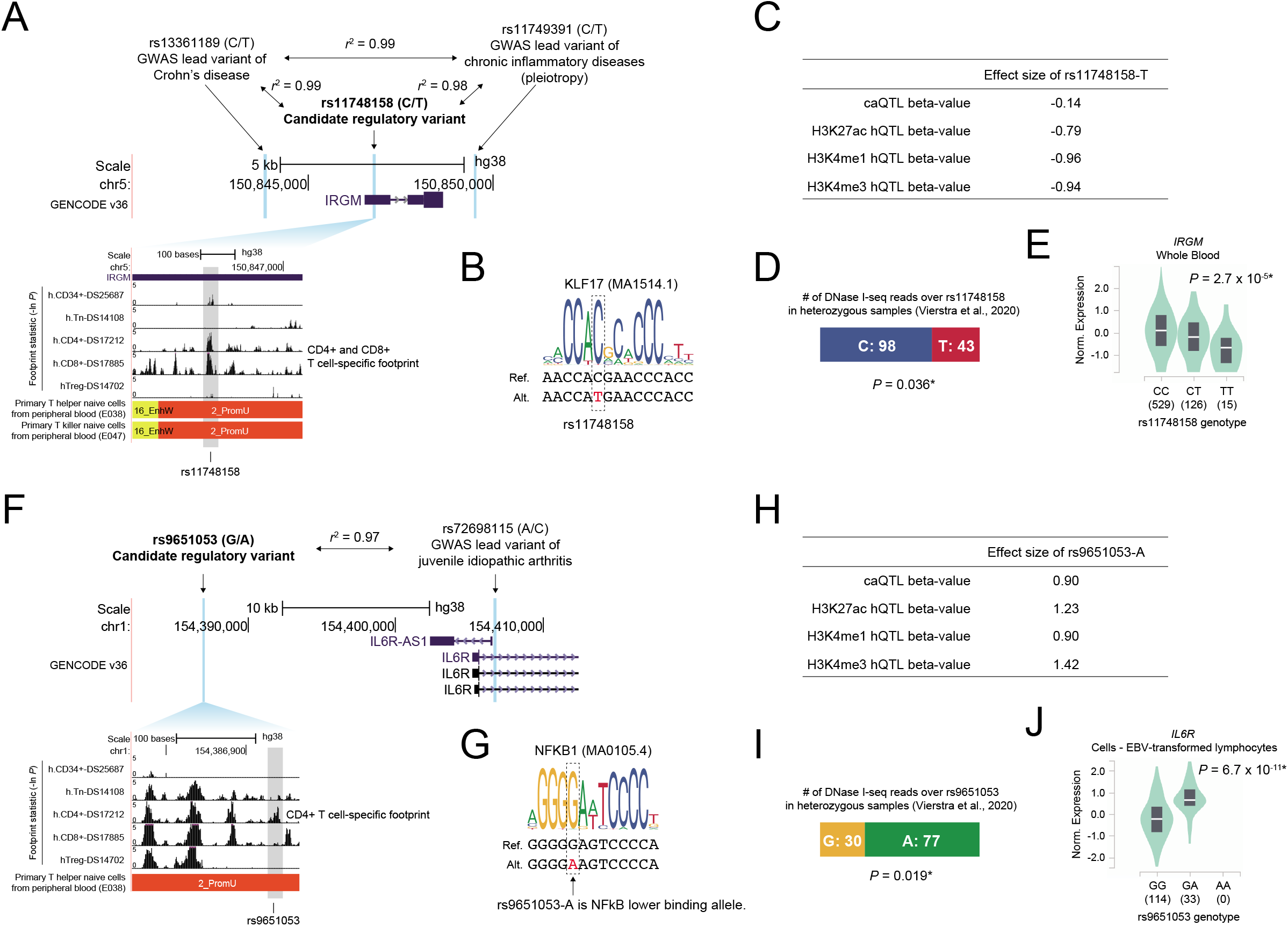
Molecular mechanisms of candidate regulatory variants. (A) The variant rs11748158 in the *IRGM* region and the magnified view with TF footprint statistics and chromatin states. Vertical blue bars indicate the location of a candidate regulatory variant and GWAS lead variants (rs13361189 and rs11749391). (B) Sequence logos based on the position weight matrix of binding sites for KLF17 (MA1514.1) are represented with genomic sequences. The location of rs11748158 is indicated by a dotted box. (C) Table of the effect sizes of rs11748158-T on chromatin accessibility (caQTL) and histone modifications (H3K27ac, H3K4me1, and H3K4me3 hQTL). (D) Number of DNase I-seq reads over each allele of rs11748158 (C/T) in heterozygous samples.^15^ A *P*-value calculated using the beta-binomial test is shown. The asterisk indicates statistical significance (*P* < 0.05). (E) Comparison of *IRGM* expression levels among rs11748158 genotypes in whole blood. The plot was generated by the GTEx project. The x-axis shows genotypes, with the number of individuals in parentheses. The asterisk indicates statistical significance. (F) The variant rs9651053 in the *IL6R* region and the magnified view with TF footprint statistics and chromatin states. Vertical blue bars indicate the location of a candidate regulatory variant and a GWAS lead variant (rs72698115). (G) The sequence logos based on the position weight matrix of binding sites for NFKB1 (MA0105.4) are represented with genomic sequences. The location of rs9651053 is indicated by a dotted box. (H) Table of effect sizes of rs9651053-A on chromatin accessibility (caQTL) and histone modifications (H3K27ac, H3K4me1, and H3K4me3 hQTL). (I) Number of DNase I-seq reads over each allele of rs9651053 (G/A) in heterozygous samples.^15^ A *P*-value calculated using the beta-binomial test is shown. The asterisk indicates statistical significance (*P* < 0.05). (J) Comparison of *IL6R* expression levels among rs9651053 genotypes in lymphoblastoid cells. The plot was generated by the GTEx project. The x-axis shows genotypes, with the number of individuals in parentheses. The asterisk indicates statistical significance.

We also identified rs9651053 (G/A) in an intergenic region as a candidate regulatory variant. This variant was highly linked with rs72698115 (A/C), a GWAS lead variant of juvenile idiopathic arthritis^41^ (*r*^2^ = 0.97), which is located within the *IL6R* gene region (Figure 4F). This regulatory variant was present within the helper T cell-specific TF footprint (h.CD4+-DS17212), the chromatin state of which is classified as promoter upstream TSS (2_PromU). Its minor allele (rs9651053-A, allele frequency = 0.0974 in the European population, 1000 Genomes) was predicted to result in decreased binding of NFKB1 (Figure 4G) and increase chromatin accessibility as well as the levels of H3K27ac, H3K4me1, and H3K4me3 (Figure 4H). Indeed, rs9651053 has been reported as an NFκB bQTL, and rs9651053-A was an NFκB lower binding allele. Referring to the data of DNase I-seq allelic imbalance, significantly more rs9651053-A allelic reads were mapped (*P* = 0.019, beta-binomial test) in rs9650153 heterozygous samples (Figure 4I), indicating that rs9651053-A increases the regulatory activity. The GTEx eQTL data showed that the rs9651053-A allele significantly increased *IL6R* expression in lymphoblastoid cells (*P* = 6.7 × 10^−11^) (Figure 4J). A recent study has suggested that rs9651053 is a causal variant in the risk of juvenile idiopathic arthritis.^42^ These findings provide evidence that NFκB acts as a repressor in the distal enhancer of the *IL6R* gene and rs9651053-A decreases the NFκB binding level, leading to increased enhancer activity and *IL6R* expression in helper T cells, resulting in a risk of juvenile idiopathic arthritis.

### Novel regulatory variants

We identified rs4017458 as a novel candidate regulatory variant highly linked with rs1333062 (G/T), a pleiotropic GWAS lead variant of chronic inflammatory diseases^38^ (*r*^2^ = 0.83), and with rs2274910 (G/T), a GWAS lead variant of Crohn’s disease^43^ (*r*^2^ = 0.83), which are mapped to the *ITLN1* gene region (Figure 5A). This candidate regulatory variant was present within the HSC and monocyte-specific TF footprint (h.CD34+-DS12734 and h.CD14+-DS17215), which showed an enhancer state (17_EnhW2 in HSCs and 14_EnhA2 in monocytes), whereas the GWAS lead variants were not found in TF footprints (Figure 5A). Its minor allele (rs4017458-G, allele frequency = 0.3270 in the European population, 1000 Genomes) was predicted to result in increased binding of ETS2 (Figure 5B) and increase H3K27ac and H3K4me3 levels (Figure 5C). Referring to the data on DNase I-seq allelic imbalance, significantly more rs4017458-G allelic reads were mapped (*P* = 0.0, beta-binomial test) in rs4017458 heterozygous samples (Figure 5D). GTEx eQTL data showed that the rs4017458-G allele significantly increased *CD244* expression in whole blood (*P* = 9.1 × 10^−16^) (Figure 5E), but not *ITLN1* expression (Figure S3B-D). *CD244* encodes a transmembrane receptor that controls immune response and is expressed in various types of immune cells, including monocytes.^44^ Our findings suggest that rs4017458-G strengthens ETS2 binding to the *CD244* enhancer and leads to upregulation of *CD244* expression, mainly in monocytes, resulting in a risk of chronic inflammatory diseases, including Crohn’s disease.

**Figure 5.**
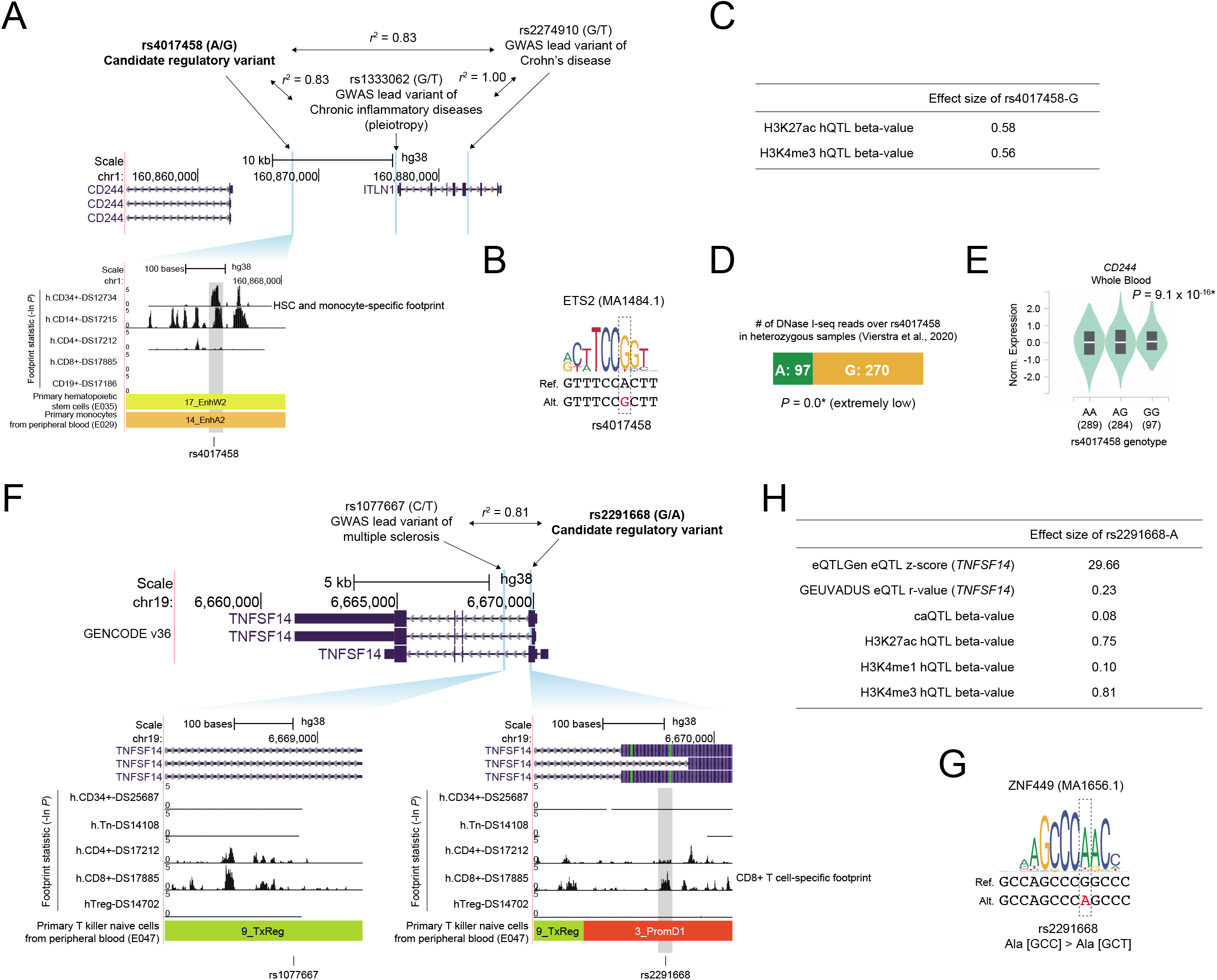
Functional impacts of novel regulatory variants. (A) The variant rs4017458 in the *CD244*-*ITLN1* region and the magnified view with TF footprint statistics and chromatin states. Vertical blue bars indicate the location of a candidate regulatory variant and GWAS lead variants (rs1333062 and rs2274910). (B) The sequence logos based on the position weight matrix of binding sites for ETS2 (MA1484.1) are represented with genomic sequences. The location of rs4017458 is indicated by a dotted box. (C) Table of effect sizes of rs4017458-G on histone modifications (H3K27ac and H3K4me3 hQTL). (D) Number of DNase I-seq reads over each allele of rs4017458 (A/G) in heterozygous samples.^15^ A *P*-value calculated using the beta-binomial test is shown. The asterisk indicates statistical significance (*P* < 0.05). (E) Comparison of *CD244* expression levels among rs4017458 genotypes in the whole blood. The plot was generated by the GTEx project. The x-axis shows genotypes, with the number of individuals in parentheses. The asterisk indicates statistical significance. (F) The variants rs2291668 and rs1077667 (GWAS lead variant) in the *TNFSF14* region and a magnified view with TF footprint statistics and chromatin states. Vertical blue bars indicate the location of each variant. (G) The sequence logos based on the position weight matrix of binding sites for ZNF449 (MA1656.1) are represented with genomic sequences. The location of rs2291668 is indicated by a dotted box. (H) Table of effect sizes of rs2291668-A on *TNFSF14* expression (eQTLGen and GEUVADIS), chromatin accessibility (caQTL) and histone modifications (H3K27ac, H3K4me1, and H3K4me3 hQTL).

We also found a novel regulatory variant, rs2291668 (G/A), located within the protein-coding region of *TNFSF14*. This variant was highly linked with rs1077667 (C/T), a GWAS lead variant of multiple sclerosis^45–47^ (*r*^2^ = 0.81), which is mapped to the intronic region of *TNFSF14* (Figure 5F). The GWAS variant was not present in a TF footprint and appeared not to affect TF binding, but the candidate regulatory variant was present in a killer T cell-specific TF footprint (h.CD8+-DS17885). The chromatin state of this region was classified as promoter downstream TSS (3_PromD1). This regulatory variant’s minor allele (rs2291668-A, allele frequency = 0.1646 in the European population, 1000 Genomes) did not change the protein sequence (Ala [GCC] > Ala [GCT]) but appeared to increase the strength of the ZNF449 binding motif (Figure 5G). In the xQTL datasets, rs2291668-A was associated with an increase in the levels of chromatin accessibility, active histone marks, and *TNFSF14* expression (Figure 5H). In a previous study,^46^ rs1077667 was predicted to be a causal variant by genotype imputation and fine mapping, but our results indicated that rs2291668 was one of the true causal variants. The variant rs2291668 is present in a protein-coding exon and is a synonymous variant, so its potential as a regulatory variant might have been overlooked thus far. TNFSF14, also known as LIGHT or CD258, is a member of the tumor necrosis factor (TNF) ligand superfamily, and its role in the pathogenesis of autoimmune diseases has been well studied.^48–51^ Altogether, these results suggest that rs2291668-A promotes ZNF449 binding to the *TNFSF14* coding exon and leads to upregulation of *TNFSF14* expression in killer T cells, resulting in the risk of multiple sclerosis. This example emphasizes the importance of the function of synonymous variants in gene regulation and disease risk.

## Discussion

In this work, we developed a computational framework that we used to discover functional variants in the TF footprints of 47 hematopoietic cells by integrating a variety of QTL datasets. We identified candidate regulatory variants highly linked to the GWAS lead variants of immune system diseases, including several common autoimmune diseases. The regulatory variants were found to be enriched in promoters and enhancers that are active in hematopoietic cells and immune tissues. The analysis identified trait–cell and trait–TF enrichment patterns, which suggested the presence of distinct pathological pathways. We found credible regulatory variants that had already been mentioned in previous studies and some that were newly identified in this study, including a variant in a protein-coding exon that does not affect the protein sequence but might alter TF recognition. Unlike previous studies, which relied on low-resolution data, such as ChIP-seq results, the use of TF footprint data from various hematopoietic cell types enabled us to precisely identify the responsible cell types and TFs in this study. Although these results are predictions and functional validation using cells or mouse models will be needed, we have provided an extensive set of possible functional variants, which provide new insights into the molecular pathogenesis of immune system diseases.

We found interesting trait–cell and trait–TF associations involved in immune system diseases. By connecting the three layers—traits, cell, and TFs—we will be able to record detailed structures of biological pathways for each trait. We have not yet been able to analyze these data because there are a limited number of variants. In this study, we focused only on immune-related traits, but this approach could be extended, using TF footprints datasets of the 243 cell and tissue types provided by ENCODE,^15^ to further discover novel aspects of a range of complex traits.

We used multiple QTL datasets (eQTL, caQTL, bQTL, and hQTL) to search for regulatory variants. However, other types of QTL have also been generated recently, e.g., splicing QTL (sQTL),^52^ DNA methylation QTL (meQTL),^53^ protein QTL (pQTL),^54^ transcription initiation QTL (tiQTL), directionality of initiation QTL (diQTL),^55^ promoter interacting eQTL (pieQTL),^56^ and mRNA *N*^6^- methyladenosine QTL (m6A QTL).^57^ TiQTL and diQTL are variants associated with enhancer RNA expression levels, and pieQTL is eQTL overlapping active regulatory elements that interact with their target gene promoters. By integrating these types of QTL, we will be able to find a larger number of functional variants in the human genome.

We discovered a regulatory variant linked with the risk of multiple sclerosis in a protein-coding exon of the *TNFSF14* gene. The regulatory functions of protein-coding exons have been identified previously; approximately 15% of the codons of the human genome are considered to serve as TF binding sites,^58^ and some exons serve as enhancers of nearby genes.^59–61^ Although we have described one case in this study, the whole picture of the roles of synonymous and nonsynonymous genomic variants within protein-coding exons in gene regulation remains unknown. TF footprinting using DNase I-seq data has enabled us to determine the precise location of TF binding sites within the chromatin with nucleotide-level resolution, so we can detect novel deleterious variants that alter TF binding levels within protein-coding exons. Although we focused on genomic variants existing in the human population, our results also suggest the presence of overlooked somatic mutations that impact gene regulation within protein-coding sequences. Recurrent mutations in protein-coding exons detected by whole-genome sequencing or exome sequencing of tumor samples have been considered to affect the structure and function of the protein encoded by a gene. However, as discussed here, some mutations in protein-coding exons could disrupt existing TF binding sites or create new TF binding sites, resulting in dysregulation of transcription. Therefore, in the future, integrated analysis of TF footprints and somatic mutation data from tumor samples will enable us to discover novel deleterious mutations in protein-coding regions affecting gene expression.

The vast majority of functional variants in the human genome still remain unexplored. Our results illustrate that an integrated analysis of TF footprints and xQTL is valuable for identifying the details of functional variants and their roles in the mechanisms of diseases. Further computational work with experimental validation will help in the identification of clinically relevant variants, leading to a better understanding of molecular pathogenesis and the discovery of novel markers and therapeutic targets.

## Supplemental Data

Supplemental Data include three figures and two tables.

## Acknowledgements

We thank all members of Mikita Suyama’s laboratory for their valuable discussions. This work was supported by a Grant-in-Aid for JSPS Research Fellow to NK (JP18J11775) from the Japan Society for the Promotion of Science.

## Data and code availability

All data used in this study are publicly available. A file of GWAS lead variants associated with “immune system disease” is available at https://www.ebi.ac.uk/gwas/efotraits/EFO_0000540, and summary files used in this study are available at https://www.ebi.ac.uk/gwas/docs/file-downloads, with the file names “gwas_catalog_v1.0.2-associations_e100_r2020-06-17_population.tsv” and “gwas_catalog-ancestry_r2020-06-17.tsv.” The eQTL datasets are available at https://storage.googleapis.com/gtex_analysis_v8/single_tissue_qtl_data/GTEx_Analysis_v8_eQTL.tar for GTEx, http://www.ebi.ac.uk/arrayexpress/files/E-GEUV-1/EUR373.gene.cis.FDR5.best.rs137.txt.gz for GEUVADIS, https://www.eqtlgen.org/cis-eqtls.html for eQTLGen, http://ftp.ebi.ac.uk/pub/databases/blueprint/blueprint_Epivar/qtl_as/QTL_RESULTS/ for BLUEPRINT, and https://dice-database.org/downloads for DICE. The caQTL dataset is available at https://zenodo.org/record/1405945/files/lead_caQTL_variants.tsv.gz. The bQTL datasets for JUND,

NFκB, POU2F1, PU-1, and STAT1 are available at https://ars.els-cdn.com/content/image/1-s2.0S0092867416303397-mmc2.xlsx, and the dataset for CTCF is available at https://www.ebi.ac.uk/birney-srv/CTCF-QTL/. The hQTL datasets are available at http://mitra.stanford.edu/kundaje/portal/chromovar3d/QTLs/. The TF footprints datasets are available at https://resources.altius.org/~jvierstra/projects/footprinting.2020/per.dataset/, and we obtained “interval.all.fps.0.05.bed.gz” for each biosample. The data of variants tested for the imbalance is available at https://resources.altius.org/~jvierstra/projects/footprinting.2020/allelic_imbalance/tested_snvs_padj.bed.gz. The datasets of chromatin states are available at https://egg2.wustl.edu/roadmap/data/byFileType/chromhmmSegmentations/ChmmModels/imputed12marks/jointModel/final/, and we obtained the files named “_25_imputed12marks_hg38lift_mnemonics.bed.gz” for each biosample.

## References

1. Tam, V., Patel, N., Turcotte, M., Bossé, Y., Paré, G., and Meyre, D. (2019). Benefits and limitations of genome-wide association studies. Nat. Rev. Genet. 20, 467–484.

2. Cookson, W., Liang, L., Abecasis, G., Moffatt, M., and Lathrop, M. (2009). Mapping complex disease traits with global gene expression. Nat. Rev. Genet. 10, 184–194.

3. Pai, A.A., Pritchard, J.K., and Gilad, Y. (2015). The Genetic and Mechanistic Basis for Variation in Gene Regulation. PLoS Genet. 11, e1004857.

4. The GTEx Consortium. (2020). The GTEx Consortium atlas of genetic regulatory effects across human tissues. Science 369, 1318–1330.

5. Lappalainen, T., Sammeth, M., Friedländer, M.R.,Hoen, P.A.C. ‘t, Monlong, J., Rivas, M.A., Gonzàlez-Porta, M., Kurbatova, N., Griebel, T., Ferreira, P.G., et al. (2013). Transcriptome and genome sequencing uncovers functional variation in humans. Nature 501, 506–511.

6. Võsa, U., Claringbould, A., Westra, H.-J., Bonder, M.J., Deelen, P., Zeng, B., Kirsten, H., Saha, A., Kreuzhuber, R., Kasela, S., et al. (2018). Unraveling the polygenic architecture of complex traits using blood eQTL meta-analysis. Biorxiv 447367.

7. Hormozdiari, F., Gazal, S., Geijn, B. van de, Finucane, H.K., Ju, C.J.-T., Loh, P.-R., Schoech, A., Reshef, Y., Liu, X., O’Connor, L., et al. (2018). Leveraging molecular quantitative trait loci to understand the genetic architecture of diseases and complex traits. Nat. Genet. 50, 1041–1047.

8. Schmiedel, B.J., Singh, D., Madrigal, A., Valdovino-Gonzalez, A.G., White, B.M., Zapardiel-Gonzalo, J., Ha, B., Altay, G., Greenbaum, J.A., McVicker, G., et al. (2018). Impact of genetic polymorphisms on human immune cell gene expression. Cell 175, 1701–1715.

9. Kumasaka, N., Knights, A.J., and Gaffney, D.J. (2018). High-resolution genetic mapping of putative causal interactions between regions of open chromatin. Nat. Genet. 51, 128–137.

10. Tehranchi, A.K., Myrthil, M., Martin, T., Hie, B.L., Golan, D., and Fraser, H.B. (2016). Pooled ChIP-Seq links variation in transcription factor binding to complex disease risk. Cell 165, 730–741.

11. Ding, Z., Ni, Y., Timmer, S.W., Lee, B.-K., Battenhouse, A., Louzada, S., Yang, F., Dunham, I., Crawford, G.E., Lieb, J.D., et al. (2014). Quantitative genetics of CTCF binding reveal local sequence effects and different modes of X-chromosome association. PLoS Genet. 10, e1004798.

12. Grubert, F., Zaugg, J.B., Kasowski, M., Ursu, O., Spacek, D.V., Martin, A.R., Greenside, P., Srivas, R., Phanstiel, D.H., Pekowska, A., et al. (2015). Genetic control of chromatin states in humans involves local and distal chromosomal interactions. Cell 162, 1051–1065.

13. Hesselberth, J.R., Chen, X., Zhang, Z., Sabo, P.J., Sandstrom, R., Reynolds, A.P., Thurman, R.E., Neph, S., Kuehn, M.S., Noble, W.S., et al. (2009). Global mapping of protein-DNA interactions in vivo by digital genomic footprinting. Nat. Methods 6, 283–289.

14. Vierstra, J., and Stamatoyannopoulos, J.A. (2016). Genomic footprinting. Nat. Methods 13, 213– 221.

15. Vierstra, J., Lazar, J., Sandstrom, R., Halow, J., Lee, K., Bates, D., Diegel, M., Dunn, D., Neri, F., Haugen, E., et al. (2020). Global reference mapping of human transcription factor footprints. Nature 583, 729–736.

16. MacArthur, J., Bowler, E., Cerezo, M., Gil, L., Hall, P., Hastings, E., Junkins, H., McMahon, A., Milano, A., Morales, J., et al. (2017). The new NHGRI-EBI Catalog of published genome-wide association studies (GWAS Catalog). Nucleic Acids Res. 45, D896–D901.

17. Machiela, M.J., and Chanock, S.J. (2015). LDlink: a web-based application for exploring population-specific haplotype structure and linking correlated alleles of possible functional variants. Bioinformatics 31, 3555–3557.

18. 1000 Genomes Project Consortium, Auton, A., Brooks, L.D., Durbin, R.M., Garrison, E.P., Kang, H.M., Korbel, J.O., Marchini, J.L., McCarthy, S., McVean, G.A., et al. (2015). A global reference for human genetic variation. Nature 526, 68–74.

19. Quinlan, A.R., and Hall, I.M. (2010). BEDTools: a flexible suite of utilities for comparing genomic features. Bioinformatics 26, 841–842.

20. Heinz, S., Benner, C., Spann, N., Bertolino, E., Lin, Y.C., Laslo, P., Cheng, J.X., Murre, C., Singh, H., and Glass, C.K. (2010). Simple combinations of lineage-determining transcription factors prime cis-regulatory elements required for macrophage and B cell identities. Mol. Cell 38, 576–589.

21. Ernst, J., and Kellis, M. (2017). Chromatin-state discovery and genome annotation with ChromHMM. Nat. Protoc. 12, 2478–2492.

22. Benjamini, Y., and Hochberg, Y. (1995). Controlling the false discovery rate: A practical and powerful approach to multiple testing. J. Royal Stat. Soc. Ser. B 57, 289–300.

23. Fornes, O., Castro-Mondragon, J.A., Khan, A., van der Lee, R., Zhang, X., Richmond, P.A., Modi, B.P., Correard, S., Gheorghe, M., Baranašić, D., et al. (2019). JASPAR 2020: update of the open-access database of transcription factor binding profiles. Nucleic Acids Res. 48, D87–D92.

24. Haeussler, M., Zweig, A.S., Tyner, C., Speir, M.L., Rosenbloom, K.R., Raney, B.J., Lee, C.M., Lee, B.T., Hinrichs, A.S., Gonzalez, J.N., et al. (2019). The UCSC Genome Browser database: 2019 update. Nucleic Acids Res. 47, D853–D858.

25. Farh, K.K.-H., Marson, A., Zhu, J., Kleinewietfeld, M., Housley, W.J., Beik, S., Shoresh, N., Whitton, H., Ryan, R.J.H., Shishkin, A.A., et al. (2015). Genetic and epigenetic fine mapping of causal autoimmune disease variants. Nature 518, 337–343.

26. Tsoi, L.C., Stuart, P.E., Tian, C., Gudjonsson, J.E., Das, S., Zawistowski, M., Ellinghaus, E., Barker, J.N., Chandran, V., Dand, N., et al. (2017). Large scale meta-analysis characterizes genetic architecture for common psoriasis associated variants. Nat. Commun. 8, 15382.

27. Georgolopoulos, G., Iwata, M., Psatha, N., Nishida, A., Som, T., Yiangou, M., Stamatoyannopoulos, J.A., and Vierstra, J. (2020). Chromatin dynamics during hematopoiesis reveal discrete regulatory modules instructing differentiation. Biorxiv 2020.04.02.022566.

28. Kikushige, Y., Ishikawa, F., Miyamoto, T., Shima, T., Urata, S., Yoshimoto, G., Mori, Y., Iino, T., Yamauchi, T., Eto, T., et al. (2011). Self-renewing hematopoietic stem cell is the primary target in pathogenesis of human chronic lymphocytic leukemia. Cancer Cell 20, 246–259.

29. Jin, Y., Chen, K., Paepe, A.D., Hellqvist, E., Krstic, A.D., Metang, L., Gustafsson, C., Davis, R.E., Levy, Y.M., Surapaneni, R., et al. (2018). Active enhancer and chromatin accessibility landscapes chart the regulatory network of primary multiple myeloma. Blood 131, 2138–2150.

30. Gruber, T.A., Larson Gedman, A., Zhang, J., Koss, C.S., Marada, S., Ta, H.Q., Chen, S.-C., Su, X., Ogden, S.K., Dang, J., et al. (2012). An Inv(16)(p13.3q24.3)-encoded CBFA2T3-GLIS2 fusion protein defines an aggressive subtype of pediatric acute megakaryoblastic leukemia. Cancer Cell 22, 683–697.

31. Masetti, R., Pigazzi, M., Togni, M., Astolfi, A., Indio, V., Manara, E., Casadio, R., Pession, A., Basso, G., and Locatelli, F. (2013). CBFA2T3-GLIS2 fusion transcript is a novel common feature in pediatric, cytogenetically normal AML, not restricted to FAB M7 subtype. Blood 121, 3469–3472.

32. Shima, H., Takamatsu-Ichihara, E., Shino, M., Yamagata, K., Katsumoto, T., Aikawa, Y., Fujita, S., Koseki, H., and Kitabayashi, I. (2018). Ring1A and Ring1B inhibit expression of Glis2 to maintain murine MOZ-TIF2 AML stem cells. Blood 131, 1833–1845.

33. Li, S., Miao, T., Sebastian, M., Bhullar, P., Ghaffari, E., Liu, M., Symonds, A.L.J., and Wang, P. (2012). The transcription factors Egr2 and Egr3 are essential for the control of inflammation and antigen-induced proliferation of B and T Cells. Immunity 37, 685–696.

34. Morita, K., Okamura, T., Inoue, M., Komai, T., Teruya, S., Iwasaki, Y., Sumitomo, S., Shoda, H., Yamamoto, K., and Fujio, K. (2016). Egr2 and Egr3 in regulatory T cells cooperatively control systemic autoimmunity through Ltbp3-mediated TGF-β3 production. Proc. National Acad. Sci. 113, E8131–E8140.

35. Ohshima, N., Takahashi, M., and Hirose, F. (2003). Identification of a human homologue of the DREF transcription factor with a potential role in regulation of the histone H1 gene. J. Biol. Chem. 278, 22928–22938.

36. Jin, Y., Li, R., Zhang, Z., Ren, J., Song, X., and Zhang, G. (2020). ZBED1/DREF: A transcription factor that regulates cell proliferation. Oncol. Lett. 20, 137.

37. Parkes, M., Barrett, J.C., Prescott, N.J., Tremelling, M., Anderson, C.A., Fisher, S.A., Roberts, R.G., Nimmo, E.R., Cummings, F.R., Soars, D., et al. (2007). Sequence variants in the autophagy gene IRGM and multiple other replicating loci contribute to Crohn’s disease susceptibility. Nat. Genet. 39, 830–832.

38. Ellinghaus, D., Jostins, L., Spain, S.L., Cortes, A., Bethune, J., Han, B., Park, Y.R., Raychaudhuri, S., Pouget, J.G., Hübenthal, M., et al. (2016). Analysis of five chronic inflammatory diseases identifies 27 new associations and highlights disease-specific patterns at shared loci. Nat. Genet. 48, 510–518.

39. Mehto, S., Jena, K.K., Nath, P., Chauhan, S., Kolapalli, S.P., Das, S.K., Sahoo, P.K., Jain, A., Taylor, G.A., and Chauhan, S. (2019). The Crohn’s disease risk factor IRGM limits NLRP3 inflammasome activation by impeding its assembly and by mediating its selective autophagy. Mol. Cell 73, 429–445.

40. Zhou, J., Theesfeld, C.L., Yao, K., Chen, K.M., Wong, A.K., and Troyanskaya, O.G. (2018). Deep learning sequence-based ab initio prediction of variant effects on expression and disease risk. Nat. Genet. 50, 1171–1179.

41. Hinks, A., Cobb, J., Marion, M.C., Prahalad, S., Sudman, M., Bowes, J., Martin, P., Comeau, M.E., Sajuthi, S., Andrews, R., et al. (2013). Dense genotyping of immune-related disease regions identifies 14 new susceptibility loci for juvenile idiopathic arthritis. Nat. Genet. 45, 664–669.

42. Tehranchi, A., Hie, B., Dacre, M., Kaplow, I., Pettie, K., Combs, P., and Fraser, H.B. (2019). Fine-mapping cis-regulatory variants in diverse human populations. Elife 8, e39595.

43. Barrett, J.C., Hansoul, S., Nicolae, D.L., Cho, J.H., Duerr, R.H., Rioux, J.D., Brant, S.R., Silverberg, M.S., Taylor, K.D., Barmada, M.M., et al. (2008). Genome-wide association defines more than 30 distinct susceptibility loci for Crohn’s disease. Nat. Genet. 40, 955–962.

44. Agresta, L., Hoebe, K.H.N., and Janssen, E.M. (2018). The emerging role of CD244 signaling in immune cells of the tumor microenvironment. Front. Immunol. 9, 2809.

45. The International Multiple Sclerosis Genetics Consortium & The Wellcome Trust Case Control Consortium 2. (2011). Genetic risk and a primary role for cell-mediated immune mechanisms in multiple sclerosis. Nature 476, 214–219.

46. International Multiple Sclerosis Genetics Consortium (IMSGC). (2013). Analysis of immune-related loci identifies 48 new susceptibility variants for multiple sclerosis. Nat. Genet. 45, 1353–1360.

47. International Multiple Sclerosis Genetics Consortium (IMSGC). (2019). Multiple sclerosis genomic map implicates peripheral immune cells and microglia in susceptibility. Science 365, eaav7188.

48. Wang, J., Lo, J.C., Foster, A., Yu, P., Chen, H.M., Wang, Y., Tamada, K., Chen, L., and Fu, Y.-X. (2001). The regulation of T cell homeostasis and autoimmunity by T cell–derived LIGHT. J. Clin. Invest. 108, 1771–1780.

49. Lim, S.-G., Suk, K., and Lee, W.-H. (2013). Reverse signaling from LIGHT promotes pro-inflammatory responses in the human monocytic leukemia cell line, THP-1. Cell Immunol. 285, 10– 17.

50. Maña, P., Liñares, D., Silva, D.G., Fordham, S., Scheu, S., Pfeffer, K., Staykova, M., and Bertram, E.M. (2013). LIGHT (TNFSF14/CD258) is a decisive factor for recovery from experimental autoimmune encephalomyelitis. J. Immunol. 191, 154–163.

51. Kinchen, J., Chen, H.H., Parikh, K., Antanaviciute, A., Jagielowicz, M., Fawkner-Corbett, D., Ashley, N., Cubitt, L., Mellado-Gomez, E., Attar, M., et al. (2018). Structural remodeling of the human colonic mesenchyme in inflammatory bowel disease. Cell 175, 372–386.

52. Li, Y.I., Geijn, B. van de, Raj, A., Knowles, D.A., Petti, A.A., Golan, D., Gilad, Y., and Pritchard, J.K. (2016). RNA splicing is a primary link between genetic variation and disease. Science 352, 600– 604.

53. Hannon, E., Spiers, H., Viana, J., Pidsley, R., Burrage, J., Murphy, T.M., Troakes, C., Turecki, G., O’Donovan, M.C., Schalkwyk, L.C., et al. (2015). Methylation QTLs in the developing brain and their enrichment in schizophrenia risk loci. Nat. Neurosci. 19, 48–54.

54. Melzer, D., Perry, J.R.B., Hernandez, D., Corsi, A.-M., Stevens, K., Rafferty, I., Lauretani, F., Murray, A., Gibbs, J.R., Paolisso, G., et al. (2008). A genome-wide association study identifies protein quantitative trait loci (pQTLs). PLoS Genet. 4, e1000072.

55. Kristjánsdóttir, K., Dziubek, A., Kang, H.M., and Kwak, H. (2020). Population-scale study of eRNA transcription reveals bipartite functional enhancer architecture. Nat. Commun. 11, 5963.

56. Chandra, V., Bhattacharyya, S., Schmiedel, B.J., Madrigal, A., Gonzalez-Colin, C., Fotsing, S., Crinklaw, A., Seumois, G., Mohammadi, P., Kronenberg, M., et al. (2020). Promoter-interacting expression quantitative trait loci are enriched for functional genetic variants. Nat. Genet. 53, 110–119.

57. Zhang, Z., Luo, K., Zou, Z., Qiu, M., Tian, J., Sieh, L., Shi, H., Zou, Y., Wang, G., Morrison, J., et al. (2020). Genetic analyses support the contribution of mRNA N6-methyladenosine (m6A) modification to human disease heritability. Nat. Genet. 52, 939–949.

58. Stergachis, A.B., Haugen, E., Shafer, A., Fu, W., Vernot, B., Reynolds, A., Raubitschek, A., Ziegler, S., LeProust, E.M., Akey, J.M., et al. (2013). Exonic transcription factor binding directs codon choice and affects protein evolution. Science 342, 1367–1372.

59. Tumpel, S., Cambronero, F., Sims, C., Krumlauf, R., and Wiedemann, L.M. (2008). A regulatory module embedded in the coding region of Hoxa2 controls expression in rhombomere 2. Proc. National Acad. Sci. 105, 20077–20082.

60. Birnbaum, R.Y., Clowney, E.J., Agamy, O., Kim, M.J., Zhao, J., Yamanaka, T., Pappalardo, Z., Clarke, S.L., Wenger, A.M., Nguyen, L., et al. (2012). Coding exons function as tissue-specific enhancers of nearby genes. Genome Res. 22, 1059–1068.

61. Mercer, T.R., Edwards, S.L., Clark, M.B., Neph, S.J., Wang, H., Stergachis, A.B., John, S., Sandstrom, R., Li, G., Sandhu, K.S., et al. (2013). DNase I–hypersensitive exons colocalize with promoters and distal regulatory elements. Nat. Genet. 45, 852–859.

