## Supplemental Figure 1-3 for "Functional variants in hematopoietic transcription factor footprints and their roles in the risk of immune system diseases"

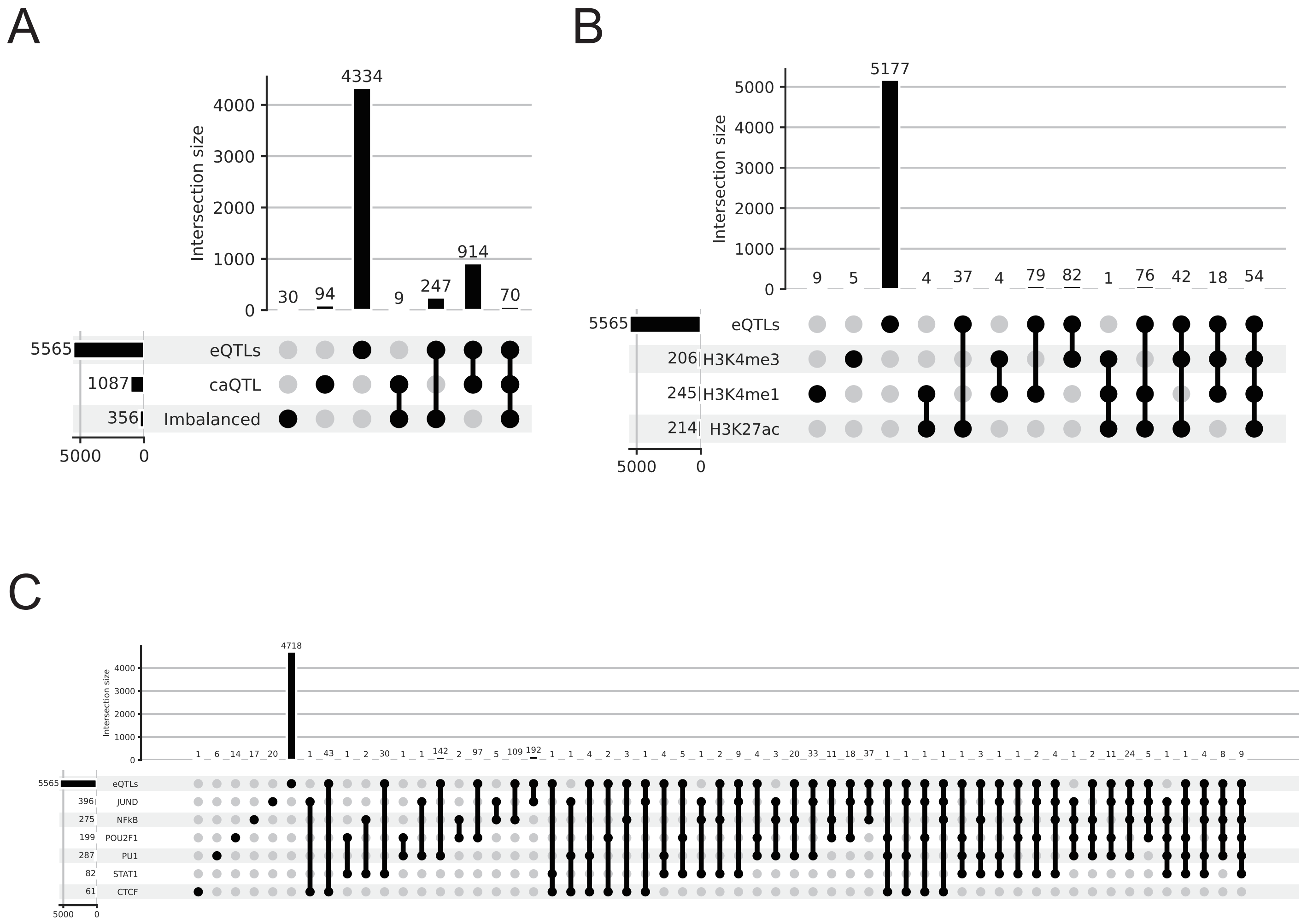


**Supplemental Figure 1. The number of variants for each combination of eQTL and other QTLs.** (A) caQTL [Kumasaka et al., 2019] and DNase I-seq allelic imbalace variants, (B) hQTL [Grubert et al., 2015], and (C) bQTL [Tehranchi et al., 2016; Ding et al., 2014].

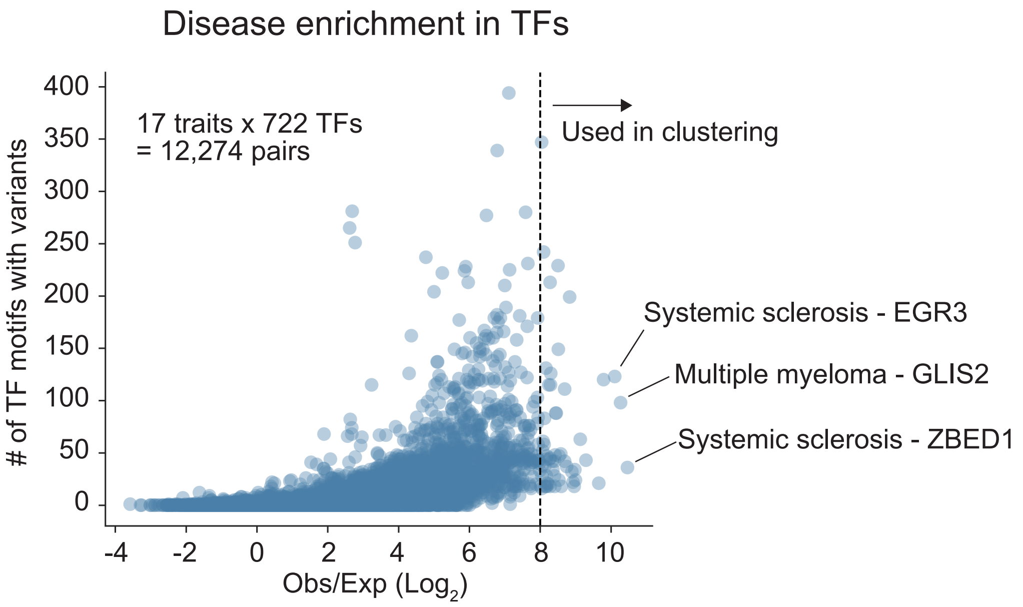


**Supplemental Figure 2. A scatter plot between the number of TF binding motifs with variants and log odds ratio.** Each point represents a trait-TF pair. TFs of which log odds ratio are higher than 8 in at least one trait were used for hierarchical clustering.

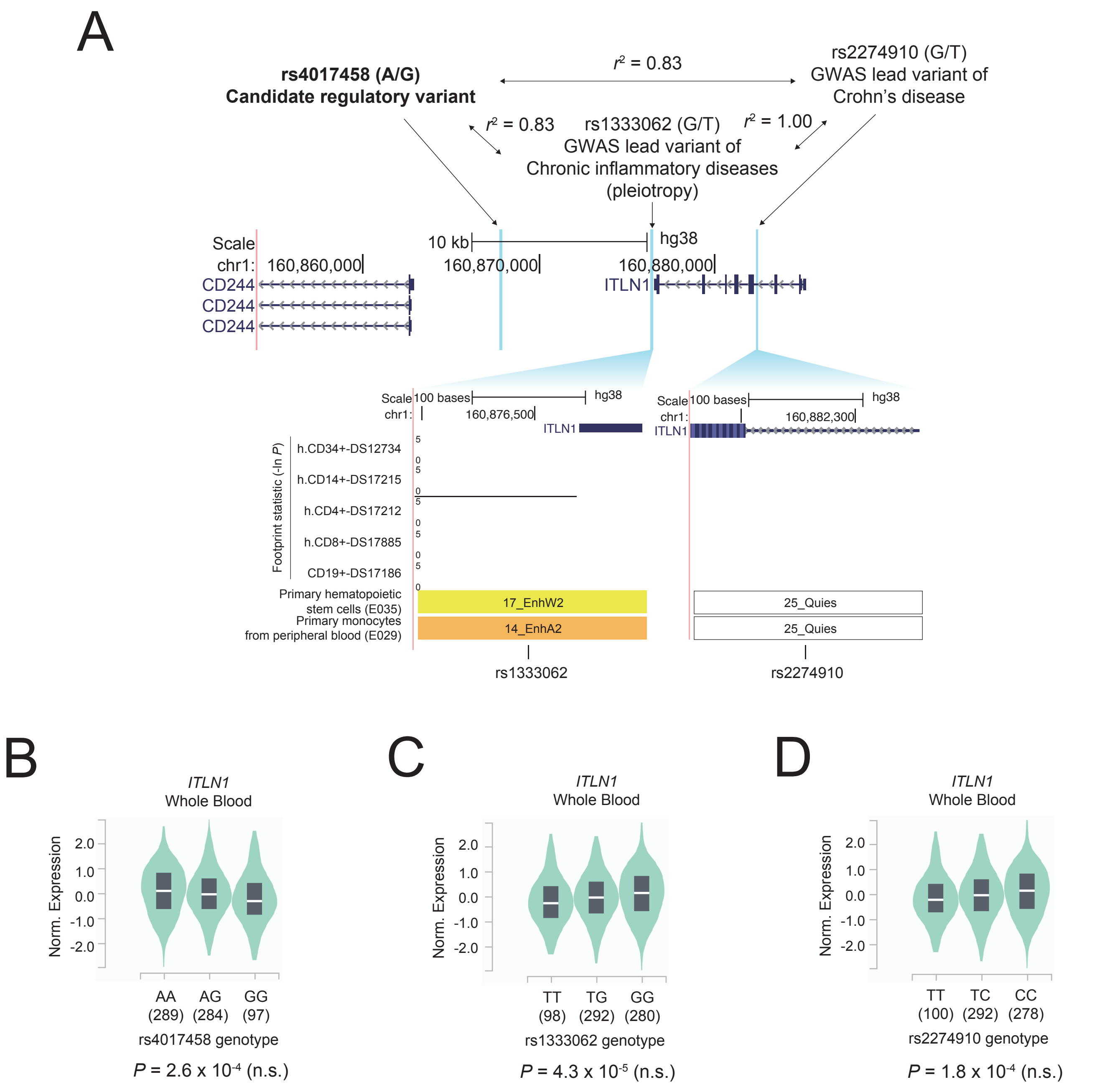


**Supplemental Figure 3. The functional impacts of GWAS lead variants linked with candidate regulatory variants rs4017458.** (A) The GWAS lead variants, rs1333062 and rs2274910, in the *ITLN1* region and the magnified view with TF footprint statistic and chromatin states. Vertical blue bars indicate the location of GWAS lead variants and a candidate regulatory variant (rs4017458). (B) Comparison of *ITLN1* expression levels among rs1333062 and (C) rs2274910 genotypes in the whole blood. The plot was generated by the GTEx project. The x-axis shows genotypes with the number of individuals in parentheses. The "n.s." indicates not statistical significance. The statistical significance was ascertained by the GTEx project.
